# High-speed two-photon microscopy with adaptive line-excitation

**DOI:** 10.1101/2024.01.24.577149

**Authors:** Yunyang Li, Shu Guo, Ben Mattison, Weijian Yang

## Abstract

We present a two-photon fluorescence microscope designed for high-speed imaging of neural activity in cellular resolution. Our microscope uses a new adaptive sampling scheme with line illumination. Instead of building images pixel by pixel via scanning a diffraction-limited spot across the sample, our scheme only illuminates the regions of interest (i.e., neuronal cell bodies), and samples a large area of them in a single measurement. Such a scheme significantly increases the imaging speed and reduces the overall laser power on the brain tissue. Using this approach, we performed high-speed imaging of the neural activity of mouse cortex *in vivo*. Our method provides a new sampling strategy in laser-scanning two-photon microscopy, and will be powerful for high-throughput imaging of neural activity.

## Introduction

Two-photon laser scanning microscopy can image deep into the tissue with a high signal-to-background ratio, tight axial confinement, and low phototoxicity^1-3^. When used in conjunction with functional fluorescence indicators, two-photon microscopy provides a powerful tool to monitor neuronal signals and activity^4-7^. Conventional two-photon microscopes build the image by raster scanning a laser focus point-by-point on the sample, resulting in a tradeoff between sampling speed, spatial resolution, and field of view (FOV). When imaging a large FOV sample with fine spatial resolution, the temporal resolution could become very poor, which may not faithfully capture the neuronal signals. In the end, the imaging throughput is limited by the single-beam scanning strategy. Another limitation of conventional two-photon microscopes is the blind scanning strategy. In particular, it is highly inefficient to record dynamic signals where the sample is repeatedly imaged in time. It not only wastes time on imaging the area without useful information, but also deposits unnecessary heat (and thus induce possible damage) to the sample.

Various types of beam multiplexing techniques^8-16^ have been proposed and demonstrated to alleviate the tradeoff between the spatiotemporal resolution and FOV, and to increase the imaging throughput. In these approaches, multiple beams, each scanning a sub-FOV, are used to image the sample. The signals are then demixed and assigned to the proper pixels in the whole image. While effective, the imaging throughput is still limited by the number of beams that could be multiplexed and the demixing quality. Furthermore, as the imaging throughput increases, the overall laser power on the tissue would inevitably increase. This could induce excessive heat on the brain tissue and thus cause tissue damage^17^.

Random-access two-photon microscope realized by acousto-optic deflectors (AODs)^18-20^, could overcome the limitation of blind scanning strategy and thus reduce the heat on the brain tissue. Instead of raster scanning, AODs could rapidly steer the beam and thus select the desirable regions of interest (ROIs) such as the neuronal cell bodies to image. We term such type of sampling strategy as adaptive sampling, as the sampling location is tailored to the ROIs on the tissue. While AODs have an increased imaging speed and reduced photodamage on the tissue, they are generally expensive, sensitive to excitation wavelength and have a limited scanning angle and thus FOV. Another strategy for adaptive sampling utilizes raster scanning with a temporally-modulated laser source^21^. However, it is not compatible with beam multiplexing schemes, and thus has a limited imaging throughput.

We report a new adaptive sampling strategy for two-photon microscopes that has high imaging speed/throughput while low laser power on the brain tissue. This microscope is optimized for population calcium imaging, where the ROIs are the neuronal cell bodies. Instead of using a diffracted-limited spot, we image the sample by scanning a short excitation line. Furthermore, we spatially modulate the line pattern so only the neuronal cell bodies are excited. Our scheme significantly increases the imaging speed as it images a large portion of the tissue in a single measurement and thus reduces the total amount of measurements per frame. Furthermore, the imaging process itself essentially pre-processes the data otherwise recorded by the point-scanning approach, as it sums up the pixels within the short-excitation line, which mostly belongs to a single ROI. The number of pixels in the recording is significantly reduced which could thus reduce the storage and alleviates the demand of the required computational resources to process the data (i.e. segmentation, and temporal traces demixing and extraction). Finally, by only illuminating the ROIs, it significantly reduces the laser power on the tissue. Using this microscope, we performed high-speed calcium imaging on mouse cortex *in vivo*. Compared to the typical two-photon microscope using point scanning strategy, we increased the imaging speed by ∼5-10× and reduced the laser power on the brain by >10×. Our new adaptive sampling strategy is compatible with beam multiplexing techniques, and holds great promise to significantly enhance the imaging throughput capabilities of two-photon microscopy.

### Principle of adaptive line-excitation

In our adaptive sampling scheme, the brain tissue is illuminated with a short line which is dynamically patterned to match the local structure of the ROIs. By scanning such a spatially modulated excitation line over the tissue, we can capture the ROIs but not the background area in the image plane (Fig. 1a-c). The spatial modulation of the excitation line is realized by a digital micromirror device (DMD), loaded with a binary mask that matches the morphology of the neuronal cell bodies (i.e. ROIs) on the sample plane. The binary mask contains the spatial footprint of the ROIs, obtained through a calibration process where we image the sample plane in high resolution, followed by segmentation. The DMD is placed at a conjugate plane of the sample plane. We shape the laser (femtosecond laser at 920 nm) beam into a line and scan it across the DMD through a resonant scanner (8 kHz) and galvanometer mirror. The deflected light from the DMD is then spatially modulated to carry the pertinent information of the ROIs, which is optically relayed to the sample plane to image the brain tissue (Fig. 1d, Methods). We note that the calibration process of the high-resolution recording is equivalent to the point-by-point scanning scheme. This can be conducted in the same microscopy, by merely configuring the DMD pattern so the ON pixels forms periodic rows (Methods, Supplementary Fig. S1-S2). Compared to the conventional point-by-point blind sampling scheme, our method has two distinct features. First, we sample a larger region in a single measurement. This reduces the number of rows in the imaging and thus increase the frame rate. In our demonstration, we shape the excitation line to be ∼11.5 μm in length (1/e^2^) on the sample plane (Fig. 1e), which is similar to the diameter of the neuronal cell body in mouse brain (10∼15 μm). This allows a fast imaging speed, while maintaining cellular resolution and avoiding excessive signal crosstalk between adjacent neurons in a single measurement, as most of the measurements would contain information of only a single neuron, in a typical labeling condition. Furthermore, our method effectively pre-processes the images otherwise recorded by the blind sampling approach, as we condense the pixels, which mostly belong to a single source, to a single pixel. This not only saves the memory to storage the raw images, but also reduces the data volume and thus the computation time in the subsequent data processing (segmentation and extraction of the temporal activity traces). Signal demixing could be processed post hoc, with the prior knowledge of all the source locations. This could resolve the signal crosstalk in the scenarios when a single measurement contains information for more than one sources. Secondly, the beam modulation from the DMD enables an exclusive sampling of the neuronal cell bodies and avoids the unnecessary excitation of background regions. It could thus greatly reduce the overall laser power delivered to the brain tissue and thus the thermal damage. Though our excitation pattern is a line, it is thin (∼2.0 μm, full-width-at-half-maximum, FWHM) (Fig. 1e). With a further leverage of the temporal focusing effect^22-27^ of the DMD, which serves as a blazed grating (Methods), our excitation pattern maintains a tight axial confinement (∼16.5 μm, FWHM) (Fig. 1f).

**Figure 1.**
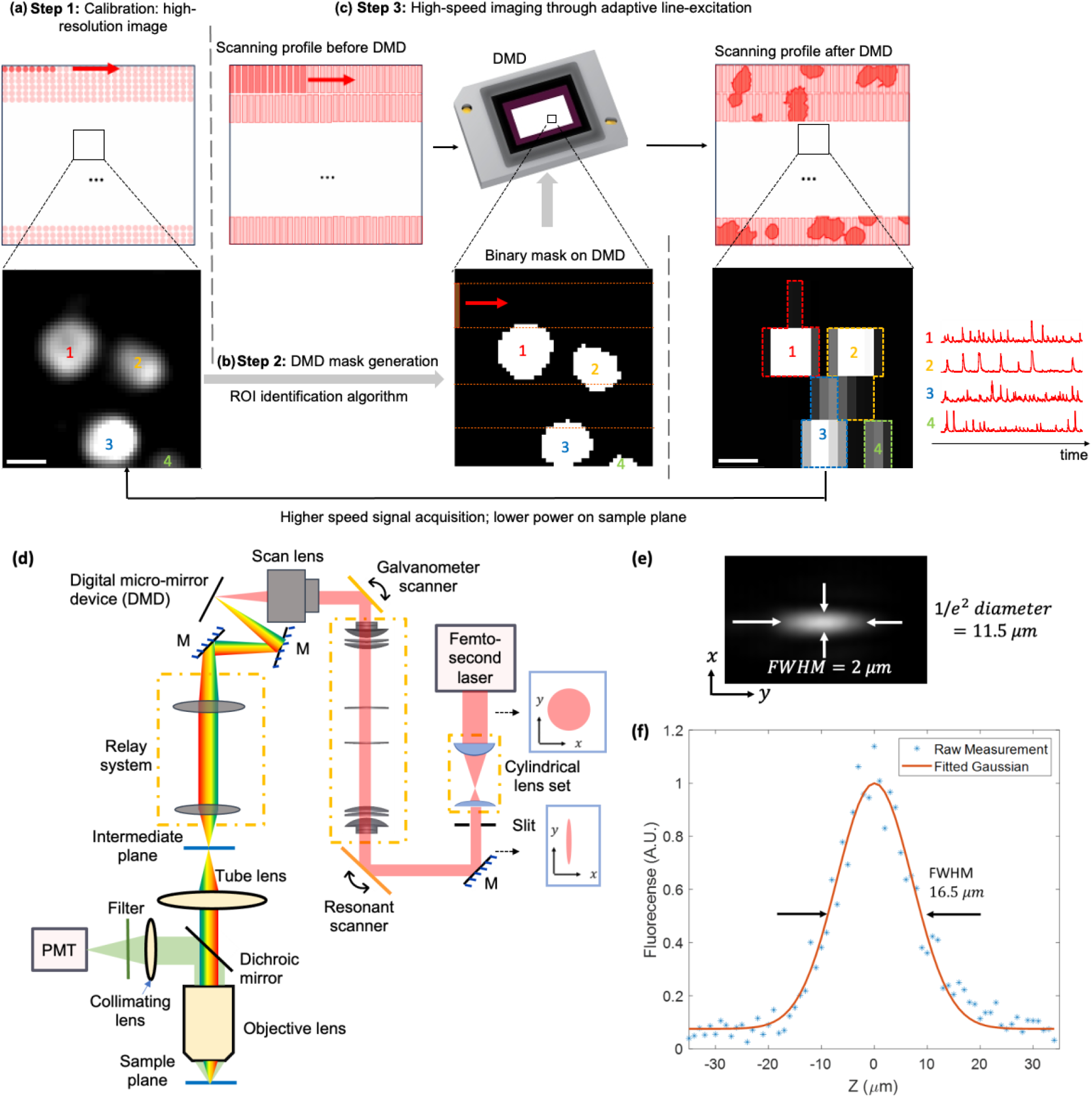
Principle, optical setup and point-spread-function (PSF) of the two-photon scanning microscope with adaptive line-excitation. (a)-(c) The working principles of adaptive line-excitation. A high-resolution video of the neuronal activity is acquired in the calibration step through an equivalent point-scanning strategy (a). The ROIs (i.e., neuronal cell bodies) could then be segmented. They are then binarized into a mask, which is loaded to the DMD (b). The laser (920 nm femtosecond laser) light is first shaped to a short line, and incident to the DMD, which is located at the conjugate plane of the sample plane and works as an intensity spatial modulator. The beam diffracted from the DMD carries the information of the ROI, and illuminates the corresponding part of the sample ROI (c). Only the ROIs but not background region are imaged. Top, illustration of the excitation scheme on the sample, with the arrow showing the beam scanning direction. Bottom, (a)(c) Zoom-in view of a sub-region of the image recorded by the photomultiplier tube (PMT); (b) Binary mask loaded on the DMD. The four ROIs are labeled in different numbers. (d) Schematic of the two-photon microscope setup with adaptive line-excitation scheme. The laser beam is first shaped into a short line, which is scanned by a resonant scanner and a galvanometer mirror onto the DMD. Light diffracted from the DMD is relayed to the sample plane through a relay system and the tube lens and objective lens. The fluorescence is detected by the PMT. M: mirror. (e) Measured PSF in the lateral direction (*xy*) for line excitation. (f) Measured axial PSF using 5 *μm* fluorescent beads. Scale bar (a), (c): 10 *μm*.

### Validation of adaptive illumination through phantom samples

We validated the beam patterning capability of the DMD and the concept of adaptive sampling through fluorescent phantom samples (Fig. 2). Using a uniform fluorescent slab, we first assessed the encoding capabilities of our system in defining arbitrary binary masks and projecting the desired patterns onto the sample plane (Fig. 2a-b). The image recorded by the PMT matched well with the binary mask loaded on the DMD, confirming the conjugate relationship between the DMD and sample plane. We then validated the binary pattern on the DMD, when projected to the sample, could indeed overlap with the ROIs on the sample structures (Fig. 2c-f). Here we used a phantom sample composed of randomly distributed fluorescent beads in diameter of 12 μm. Through a calibration process which is equivalent to point scanning (Methods, Supplementary Fig. S1-S2), we obtained a high-resolution image of the sample with 256 × 320 pixels, in a resolution of ∼2.0×2.2 μm^2^ (Fig. 2c). Such a calibration process mapped individual DMD pixels to the sample coordinates. We then transformed the high-resolution image to the DMD mask (Fig. 2d) and performed the high-speed line scanning with adaptive sampling. The acquired image matched well with the DMD mask, confirming that the mask aligned well with the sample plane (Fig. 2e). Finally, we simulated the high-speed scanning results based on the high-resolution image (Fig. 2c) and the bi-directional scanning trajectory. The simulated image (Fig. 2f) shows a high similarity with the experimental result (Fig. 2e), further verifying the robust mapping relationship between the DMD and sample structure.

**Figure 2.**
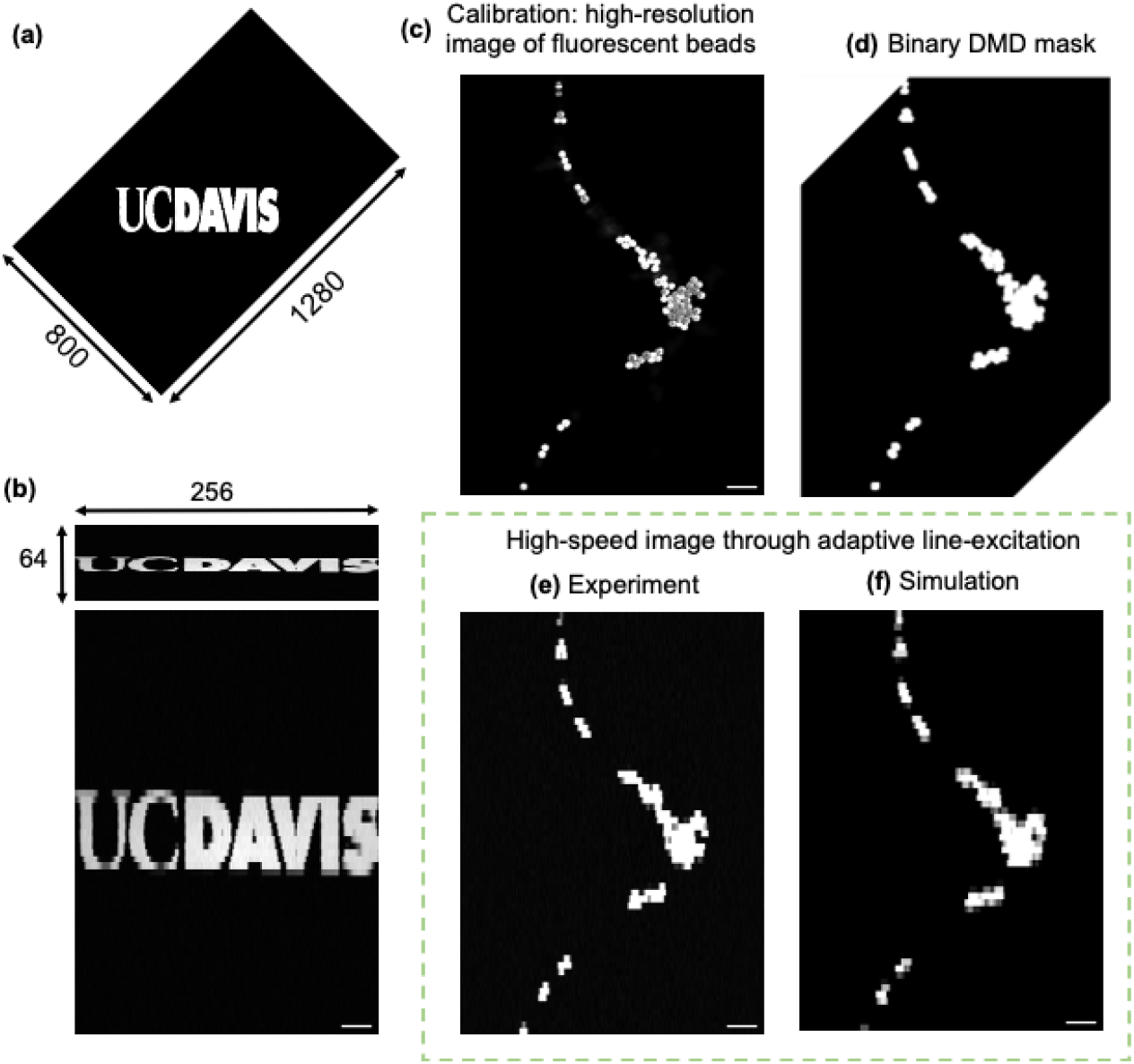
Validation of the adaptive sampling scheme through phantom samples. (a)-(b) Projection of the DMD mask onto a uniform fluorescent slab through adaptive short-line excitation. (a) DMD mask (1280×800 pixels) with characters “UC DAVIS”. (b) Top: raw image (256×64 pixels) acquired from PMT through adaptive line-excitation with bi-directional scanning; bottom: interpolated and resized image (500×695 pixels) with square pixels. (c)-(f) Imaging of a phantom sample with randomly distributed fluorescent beads, in 12 μm diameters. (c) High-resolution image of the sample acquired through an equivalent point-scanning approach. (d) Binary mask on the DMD. The two corner regions of the mask were outside the DMD active regions and were not displayed. (e) A single frame of the recording from the sample acquired in 198 Hz, using the adaptive line-excitation with bi-direction scanning. (f) Simulated high-speed image based on the binary mask and the bi-directional scanning trajectory of the resonant scanner. Scale bar: 50 *μm*.

### High-speed calcium imaging of neuronal activity in vivo

We conducted *in vivo* experiments to monitor the cortical activity of layer 2/3 in primary visual cortex (V1) in awake mouse transfected with calcium indicators GCaMP6f^28^, over a FOV of ∼500 μm × 695 μm. Following the procedure outlined in Fig. 1(a)-(c), we first calibrated the spatial location of the ROIs by obtaining the high-resolution recording of the sample plane (Fig. 3a, 256 × 320 pixels, in a resolution of 2.0 × 2.2 μm^2^, at a depth of 170 μm), and segmenting the recording using SUNS^29^ (Fig. 3c), a state-of-the-art fast segmentation algorithm on calcium imaging. In the represented example, we found 79 ROIs, which were the active neurons during the video acquisition period in calibration (∼7 mins). By loading the ROIs contours into the DMD, we conducted high-speed recording through adaptive line-excitation (Fig. 3d, 256 × 64 pixels). Notably, our sampling strategy reduced the number of rows scanned by the resonant scanner, and thus increased the frame rate. With the height of each row being ∼11 μm, and using a bi-directional scanning scheme, we achieved a frame rate of 198 Hz over the FOV ∼500 μm × 695 μm. Such a frame rate is significantly higher than the typical ones in conventional two-photon microscopes using point scanning strategy. Compared to the case where only line-excitation was used but without the adaptive sampling strategy (Fig. 3b, with the binary mask on the DMD being all 1 by turning on all DMD pixels), the adaptively sampled images (Fig. 3d) used a significantly smaller average laser power on sample (∼14× smaller than the case without adaptive sampling, as the occupied area of ROIs over the entire plane is ∼7.2%), while maintaining a similar signal-to-noise ratio. In our case of adaptive line-excitation, only 14 mW of laser power was on the brain tissue. A plain line-excitation without adaptive sampling would result in ∼194 mW, which could be near the thermal damage threshold of the mouse brain^17^. Finally, from the high-speed recording with the adaptive line-excitation strategy, we extracted the temporal activity traces of individual ROIs through CalmAn^30,31^ (Fig. 3e). As we had the pixel locations of individual ROIs as prior knowledge, we used their centroid pixels to initialize the search of the spatial footprint of individual ROIs in the algorithm. This increased the efficiency and efficacy of the algorithm.

**Figure 3.**
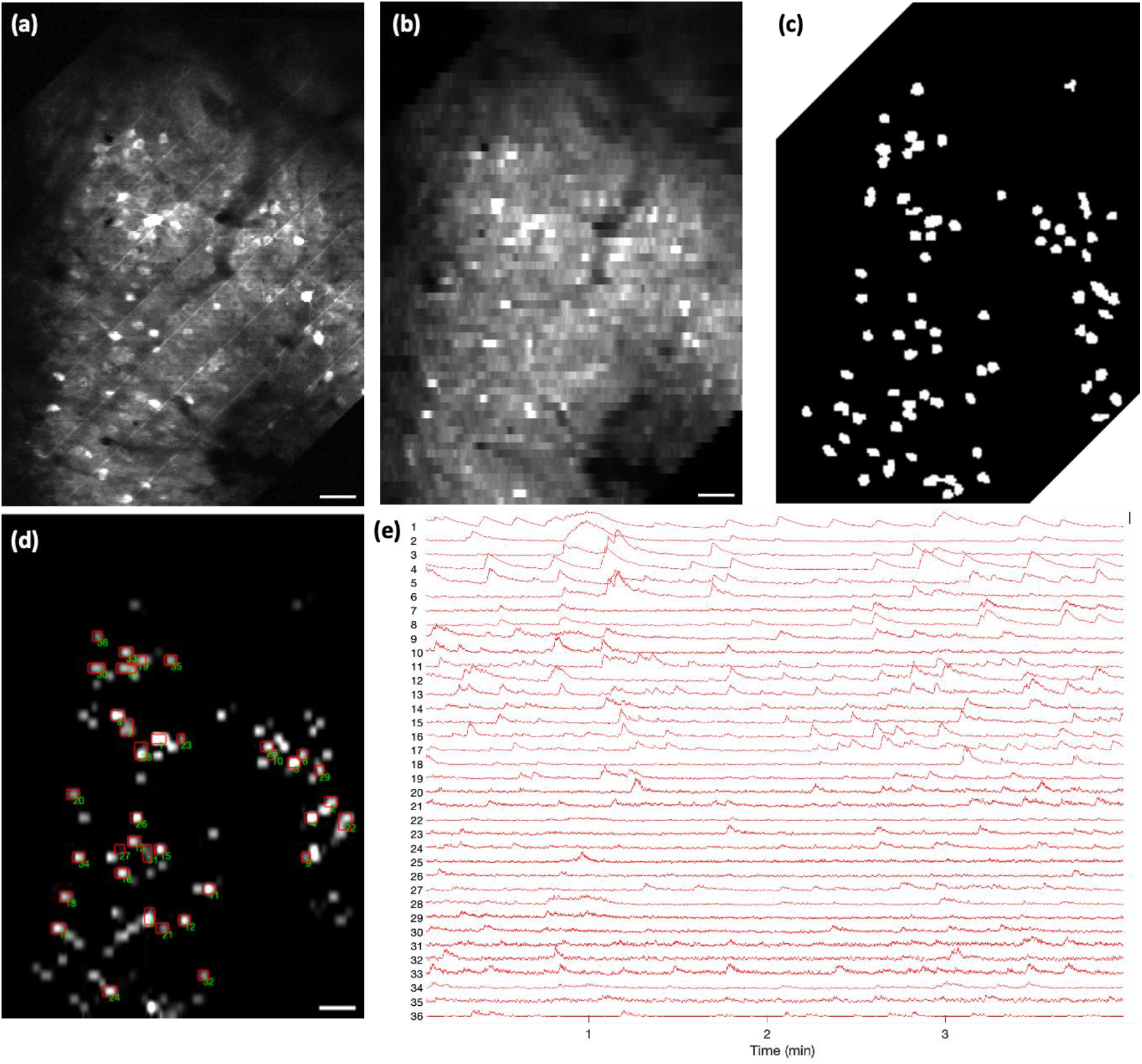
In vivo calcium imaging of mouse V1 using the adaptive line-excitation two-photon microscope. (a) Temporal averaged frame of the neuronal activity recording (256×320 pixels over 500×695 *μm*^2^ FOV) in mouse V1 at a depth of 170 μm using an equivalent point scanning method. (b) The line-scanning result of the same FOV (256×64 pixels) in (a) without adaptive sampling. (c) The binary mask for (a), constructed from the ROI segmentation algorithm. Only the neuronal cell bodies with neuronal activities during the video acquisition period in calibration (∼7 mins) were considered as active ROIs. The two corner regions of the mask were outside the DMD active regions so were not displayed. (d) The line-scanning result (256×64 pixels) with adaptive sampling strategy by applying the mask on the DMD. Only the ROIs defined on the mask in (c) were illuminated and sampled. (e) Selected temporal activity traces from the corresponding red-contoured ROIs in (d). Scale bar: (a)(b)(d) 50 *μ*m; (e) Δf/f = 2.

## Discussion

In summary, we proposed and demonstrated the concept of adaptive sampling with line illumination in two-photon microscopy. The DMD functions as the intensity spatial light modulator as well as a blazed grating for temporal focusing. By using line illumination and only exciting the regions of interest rather than the entire field of view, we could achieve a high imaging speed (up to 198 Hz with 500×695 μm^2^ FOV) while reduce the overall laser power and thus phototoxicity on the sample. Crucially, our method is compatible with many other beam multiplexing techniques, such as spatiotemporal multiplexing^9,14^, so as to further increase the imaging throughput. The capability to reject the illumination on the non-ROI background regions becomes particularly important to reduce the overall laser power and thus heat generation in the brain tissue when multiple beams are scanning in the tissue. While not demonstrated here, our method is also compatible with three-dimensional/volumetric imaging by incorporating an axial scanning mechanism (such as tunable lenses) after the DMD. While designed for high-speed imaging, our microscope retains the capability for the equivalent point-scanning high-resolution imaging, by properly setting and cycling the DMD pattern with a synchronization with the scanner, as in our calibration step.

The line-scanning approach is one type of PSF engineering to increase the imaging throughput^32^. Here, we used a short-line excitation with a single pixel detector. Two similar strategies involve pairing a long-line excitation with a camera^33^, or a long-line excitation with a linear PMT array^34^. They have challenges in imaging deep due to light scattering. Other long-line excitation strategies with a single pixel detector require imaging the sample through multiple line patterns, and reconstruct the image computationally through principles of compressive sensing^35^ or topographical imaging^36^. They typically require intense computational resources or complex system setup. Our strategy does not have these challenges. One advantage of our method is that the signal in each measurement over the short-line mostly comes from the same source. Such a strategy essentially pre-processes the data, and thus reduces the signal processing time as the number of pixels in the image is reduced. If there are multiple sources in a single short line, they could be separated through demixing algorithms such as non-negative matrix factorization (e.g. CalmAn^30,31^). We note that previous reports have demonstrated block-scanning^25,26^, where the illumination was a 5×5 μm^2^ block and their fluorescence were summed together into a single measurement. There, a low-repetition rate laser was used, and each block was sampled with one laser pulse, so as to avoid oversampling in the fast-scanning axis. The resonant scanner has to synchronize with the laser pulse clock, which required a custom-tuned resonant scanner and complicated electronic controls. Our method uses line-illumination, and the linewidth in the fast-scanning axis is thin. Thus, we could use a typical 80 MHz laser without synchronization among the scanner and laser clock. Furthermore, as the line rate is determined by the resonant frequency of the scanner, short-line scanning could achieve the same speed as block scanning.

Our adaptive sampling strategy uses a DMD to spatially modulate the excitation pattern. Compared to existing adaptive sampling approaches (AODs^18-20^ and adaptive laser source^21^), our method only requires simple hardware, and is compatible with beam multiplexing. In particular, compared with the method using a temporally modulated laser source, we do not need the synchronization of the modulator and the scanner. Like other adaptive sampling approaches, we require a prior knowledge of the ROIs, and the imaging quality is sensitive to the motion of the tissue. This could be readily resolved by including a real-time feedback loop and compensate the motion through the scanner ^20^.

In typical two-photon microscopes, there is a tradeoff among imaging throughput and required laser power on the tissue. Our adaptive line-excitation two-photon microscope alleviates this tradeoff and could simultaneously increase the imaging throughput and reduce the on-tissue laser power. It could be powerful in applications which requires high-throughput imaging, such as voltage imaging or calcium imaging over a large 3D volume.

## Methods

### System setup of the two-photon microscopy with adaptive line-excitation

We home-built the two-photon microscope with the adaptive line-excitation scheme based on an Olympus open-stand frame. The system utilizes an 80 MHz repetition rate femtosecond laser (Axon 920-2, Coherent Inc.), which has intrinsic pulse compensation and power adjustment components, producing a 920 nm-centered excitation wavelength. The beam first goes through a cylindrical lens set and a slit with adjustable width (VA100CP, Thorlabs) so its spatial profile is shaped to be a line shape. The beam is then scanned by a resonant scanner mirror (4×5 mm^2^ size, SC-30, Electro-Optical Products Corp) and a galvanometer scanner (GVS001, Thorlabs). The two scanners are conjugated to each other through a set of relay lens adopted from ref [37]. After a scan lens^37^, the beam is focused and scanned on the DMD (DLP650LNIR, Texas Instruments), which is placed at the back focal plane of the scan lens to ensure its conjugate relationship with the sample plane. After the DMD, the beam goes through a 4f relay system, followed by a tube lens (TTL200MP2, Thorlabs) and a water-immersion objective lens (CFI75 LWD 16X, Nikon). The effective magnification between the DMD plane and the sample plane is 13.3×. The excitation beam, whose pattern is dynamically modulated by the DMD as it scans, is then focused on the sample plane.

In the collection path, the fluorescent emission passes through the same objective lens, is reflected at the dichroic mirror (FF705-Di01-25x36, Semrock), and finally reaches the PMT (H7422-40, Hamamatsu) after a GFP bandpass filter (FF03-525/50-25, Semrock) and short-pass filter (FF01-715/SP-25, Semrock). The current signal from the PMT is then converted to voltage signal through a transimpedance amplifier (TIA60, Thorlabs), before being collected by a high-speed data acquisition card (vDAQ, Vidrio Technologies). ScanImage^38^ is used to control the entire system.

The lateral excitation profile of the illumination beam was characterized by imaging the exciting pattern on a uniform fluorescent slab through a camera after the slab, in a transmission configuration. The axial excitation profile of the illumination beam was characterized by taking a z-stack of a 5 μm fluorescent bead sample through a translation stage (OptiScan III, Prior).

### Configuration of the DMD in the optical system

The DMD used in our setup has 1280×800 pixels over an active region of 13.824×8.64 mm^2^. The DMD can reach a frame rate with 1-bit binary at 10.752 kHz with a development kit (SuperSpeed .65” WXGA, ViALUX). The “ON” and “OFF” states are tilted ±12° with respect to the DMD surface. It can be used as a diffraction grating^39^ with a 10.59° blazed angle under 920nm. In our system, the DMD surface is rotated by 45° along the incident beam so that the incidence beam and the diffraction beam at blazed condition are on a plane parallel to the optical table (Supplementary Fig. S3). The DMD is tilted such that the chief ray (corresponding to central wavelength of the laser light) of the diffraction beam at blazed condition propagates on axis to the subsequent optics after the DMD. The diffraction light with orders other than the order of blazed condition are filtered out before the subsequent optical components after the DMD.

To ensure the spatial focal plane of the scan lens overlaps with the temporal focal plane (i.e. the DMD surface) while the beam is being scanned, we deliberately introduced an in-plane spatial displacement (i.e. decentering^40^) of the scan lens (Supplementary Fig. S4). This introduces an aberration, which compensates for the mismatch between the spatial focusing plane and the temporal focusing plane, thereby ensuring consistent pattern of the incident line illumination across the entire surface of the DMD.

### System calibration and the generation of the high-resolution image

The adaptive sampling process begins with a calibration to acquire the high-resolution structural image of the sample. Such a calibration could be conducted by operating the microscopy in an equivalent point-scanning mode (Supplementary Fig. S1-S2). There, we designed a set of five DMD masks, each being a periodic line pattern, i.e. two out of ten lines are turned on. The lines that are turned on are shifted by two lines in the next pattern. Thus, in the first DMD mask, line numbered 1, 2, 11, 12, 21, 22, and so forth are turned as “ON” state, while the remaining lines are set to “OFF”. In the second DMD mask, lines numbered 3, 4, 13, 14, 23, 24, and so on are turned as “ON” state. This process continues until the fifth DMD mask, where lines numbered 9, 10, 19, 20, 29, 30, and so forth are turned as “ON” state. For each DMD mask, we scan the laser light in the same line-shape across the DMD and acquire an image through the PMT. Effectively, the illumination pattern becomes a spot on the sample (∼2.0×2.2 μm^2^). Through a sequential combination of the rows of the five PMT images, we could obtain one high-resolution image, equivalent to point-scanning. By cycling the five DMD masks continuously, we could record the high-resolution video continuously, in an effective frame rate of ∼21.6 Hz.

As the intensity along the excitation line is not strictly uniform, we introduce offsets for the start of the scanning position of the galvanometer mirror for each of the five DMD mask, so the center of the line illumination always overlaps with the lines that are turned as “ON” state from the each DMD mask (Supplementary Fig. S2). This ensures the illumination intensity of each spot on the high-resolution image is the same. Such a process is realized by MROI functionality in ScanImage. When the high-resolution image acquisition starts, the MROI scheme shifts the starting position of each FOV by adjusting the offset voltage, corresponding to a 2-line shift at the DMD plane. Meanwhile, a trigger signal is sent from vDAQ to the DMD, which synchronizes the DMD pattern display with MROI shifting under the slave mode.

### Data acquisition

For a resonant scanner, the angular velocity at the edge of the scan field is highly nonlinear due to the sinusoidal scanning property, and thus, the acquisition at the edge is less preferred. The ratio between the active acquisition time and the total time of a line is defined as the fill fraction in ScanImage. In the experiment, the fill fraction was 71.3% in the temporal domain and 90% in the spatial domain.

### Data processing

In the calibration process, we use SUNS^29^ to segment the neurons (i.e. ROIs) of the high-resolution recording. We manually inspected the fluorescence traces of each found ROI, and removed the ROIs whose fluorescence trace is noisy and did not show clear calcium transients. For the high-speed recorded data from the adaptive line-excitation scheme, we first denoised the data through DeepCAD^41^, a self-supervised learning algorithm. We then initialized the centroid coordinate of each ROIs through the prior knowledge of the DMD mask and scanning trajectory, in CalmAn algorithm^30,31^. The temporal activity traces of each individual ROIs could then be extracted.

### Animal Procedures

All animal experiments and housing procedures were conducted with the approval and guidance from University of California Davis Institutional Animal Care and Use Committee (IACUC). Wild-type C57BL/6J mice (2-5 months old) were injected with AAV1-hSyn-GCaMP6f into the right primary visual cortex and chronic craniotomy was performed to attach a circular 3.5 mm glass coverslip window for imaging. The coverslip center position was set to be the injection site. A custom stainless steel headplate with a 7 mm hole in the center was attached to the skull surrounding the injection and craniotomy location using dental cement (MetaBond). The coverslip was secured using cyanoacrylate (3M VetBond) and further sealed with dental cement to cover the exposed bone in the center of the headplate, and to stabilize the headplate for head-fixed imaging. After 3∼4 weeks which allows for virus expression, awake imaging was done with the headplate fixed to custom optical posts on top of a 3D printed treadmill on which the mouse was free to run.

## Acknowledgement

We acknowledge support from National Science Foundation (CAREER 1847141) and Burroughs Wellcome Fund (Career Award at the Scientific Interface: 1015761).

## Supplementary Information

**Supplementary Figure 1.**
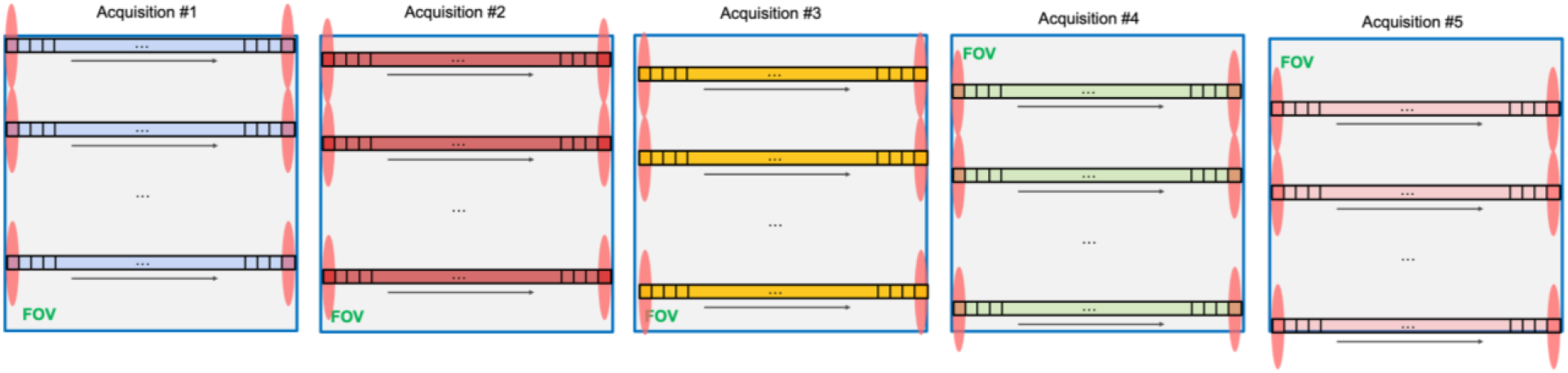
Calibration mode of the two-photon microscope, equivalent to point scanning. The high-resolution image from the equivalent point scanning is constructed from a series of 5 images from single-directional scanning. For each image, 2 lines out of 10 lines on the DMD are set as “ON” state along the single-directional scanning trajectory. The starting point of the FOV scanning is shifted deliberately to ensure the central part of excitation line shape, which has the highest intensity values of the entire line, aligns with those lines set as “ON” states in the DMD. As the DMD is located at the conjugate plane of the image plane and works as an intensity modulator, only one fifth of the area at the sample plane is illuminated. After 5 acquisitions for the 5 DMD masks, 5 PMT images are captured, and the high-resolution image is constructed by sequentially combining the active lines recorded in each image.

**Supplementary Figure 2.**
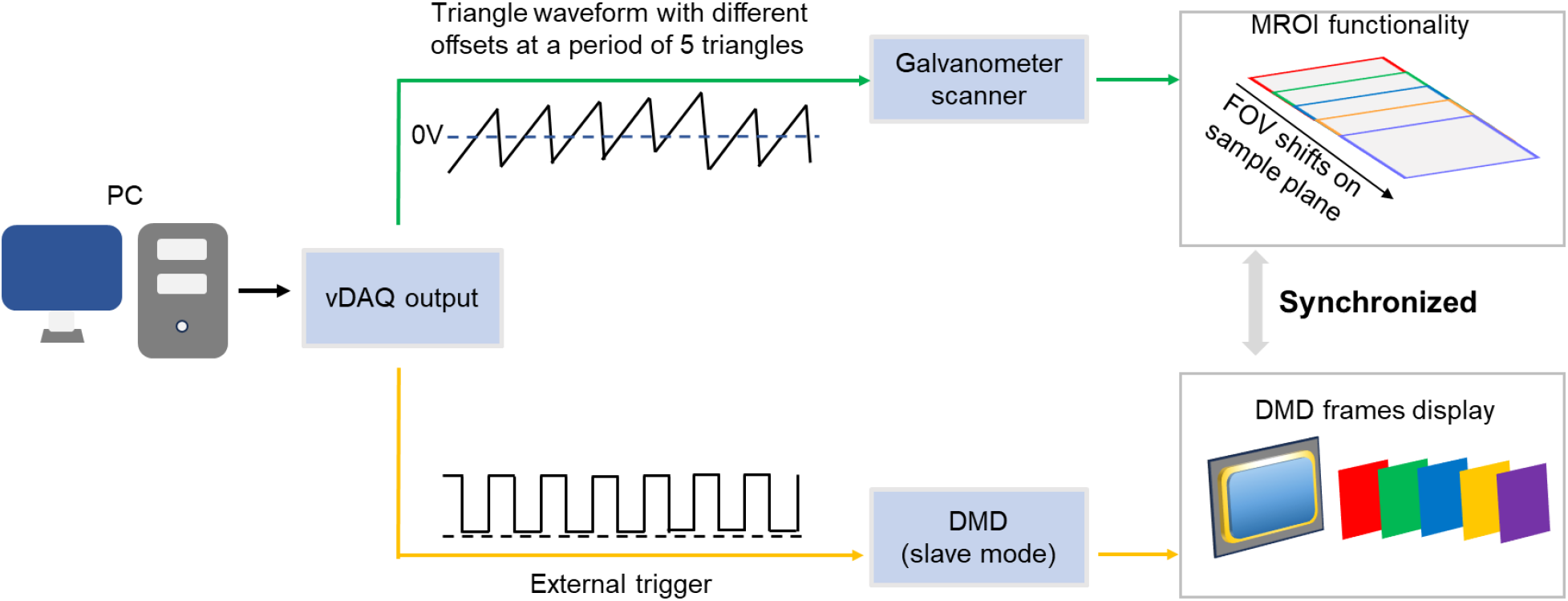
Synchronization and active MROI acquisition. To achieve the equivalent point scanning for a high-resolution image, the simultaneous shifting of the FOV in the galvanometer scanning direction and the change of DMD pattern is required. For each shift of the scanning FOV on the DMD by 2 lines, an offset voltage based on the V/deg parameter for the galvanometer scanner is applied to the command voltage from the vDAQ analog output to the galvanometer scanner. Concurrently, internal communication within vDAQ generates a digital rectangular signal, which is synchronized with the analog output voltage to the galvanometer scanner and sent to the DMD portal to update the DMD mask.

**Supplementary Figure 3.**
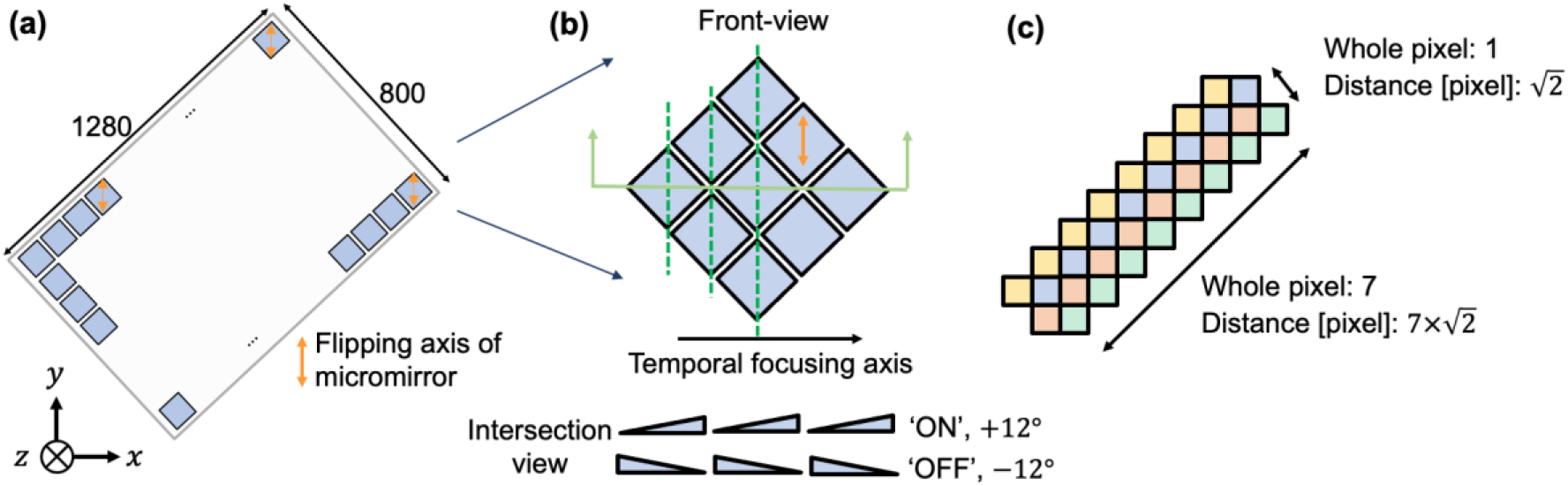
Schematics of DMD layout. (a) The front view of the whole DMD surface and (b) a zoomed-in region: the DMD surface is rotated by 45° so the hinge (i.e. flipping axis) of each micromirror is vertical to the surface of the optical table. The incident light and the diffraction light at the blazed condition are in the horizontal plane parallel with the surface of the optical table. The intersection view shows the tilting conditions of micromirrors at the “ON” and “OFF” states. (c) Adjacent DMD rows which are distinguished by colors. We define the distance between 2 micromirrors in a row as √2 pixels.

**Supplementary Figure 4:**
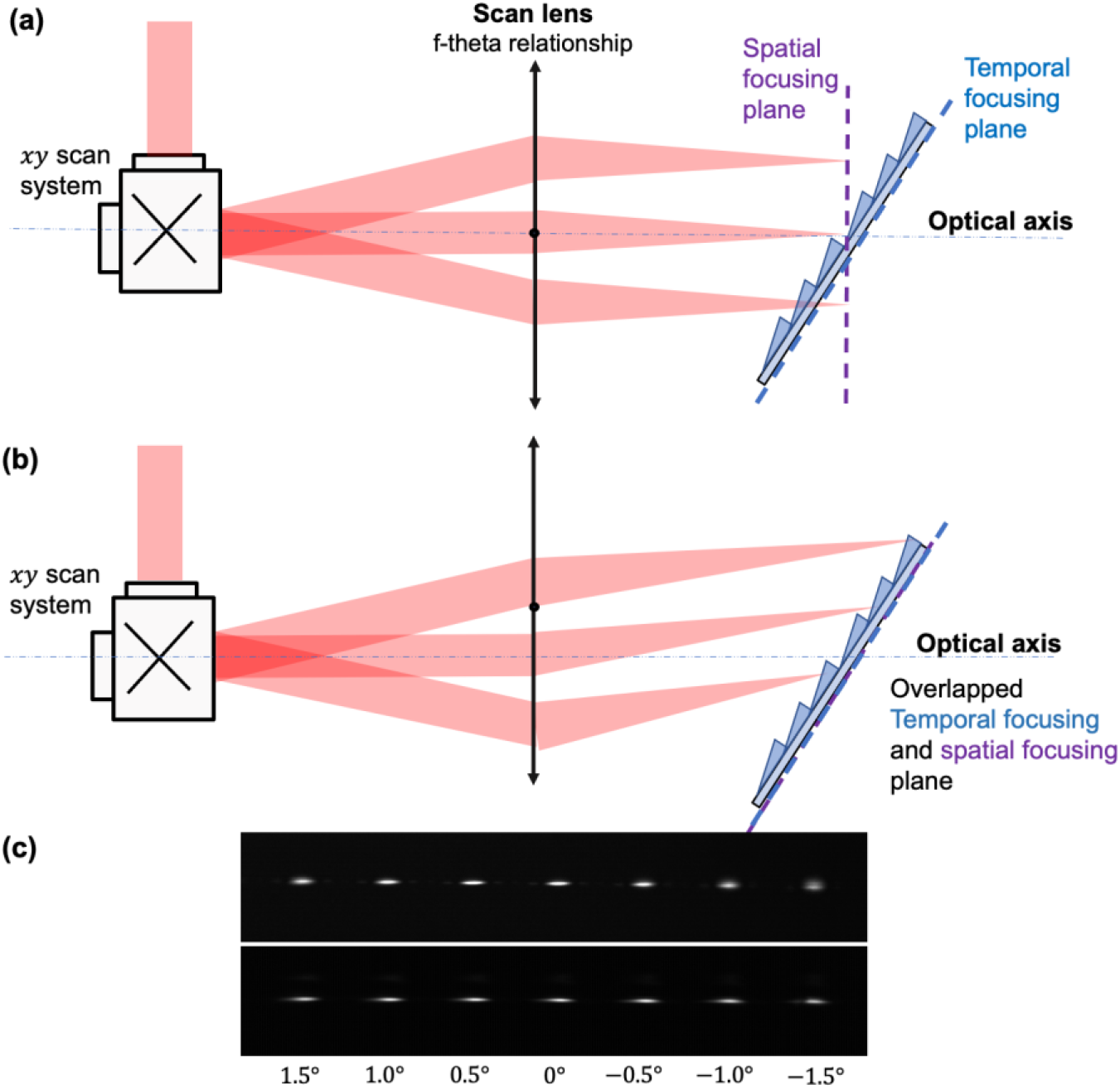
Illustration of the temporal focusing plane and spatial focusing plane. (a) The conventional scanning scenario: both the scanners and scan lens (SL) are symmetric with respect to the optical axis. Due to the f-theta relationship of the scan lens, the collimating beams from different scanning angles are focused on the plane perpendicular to the optical axis which is the focal plane of scan lens. This is different from the temporal focusing plane which is the DMD surface. (b) With a proper decentered distance between the scanner mirror and optical axis, the scanning beams from different angles will all focus on the plane of the DMD surface, which has a tilting angle with respect to the focal plane of scan lens. The temporal focusing and spatial focusing plane are then overlapped. (c) Experimentally measured focal spots on a conjugate plane of the DMD surface to compare the quality of the focal spot across the field of view (thus different scanning angle of the galvanometer mirror) between the conventional scanning scenario (top) and optimized scanning scenario (bottom). The beam profile has a more consistent shape across the field of view in the optimized scanning scenario.

